# Signatures for conserved phenolic-based acclimation responses to UV in a unicellular algal sister of land plants

**DOI:** 10.64898/2026.11.05.764526

**Authors:** Cäcilia F. Kunz, Deborah Schopp, Ilka N Abreu, Tatyana Darienko, Janine Fürst-Jansen, Stefanie König, Ivo Feussner, Maike Lorenz, Jan de Vries

## Abstract

Ultraviolet (UV) light is a major stressor that particularly comes to bear in the terrestrial habitat. As such it is among the major challenges that the earliest land plants had to overcome. The closest algal relatives of land plants, the zygnematophytes, are single celled and filamentous algae that—in the context of their terrestrial and freshwater habitats— have diverse adaptations to UV irradiance. As such, contextualized with data on land plants, zygnematophyte algae harbor important information for inferring how the earliest land plants have acclimated to UV. Here we exposed the unicellular zygnematophyte model *Mesotaenium* to daily UV irradiance and assessed its response through microscopy, untargeted metabolomics, and global gene expression profiling using transcriptomics. Our data reveal cell biological acclimation by formation of lipid droplets. Further, we find that the cells accumulated chemodiverse coumarin-like derivatives. Altogether, our data suggest that acclimation to long-term UV exposure is composed of building blocks of a molecular chassis conserved with that of land plants, underpinned by phenolics and genes that are tied to their biosynthesis. From this chassis, divergent evolutionary trajectories have assembled them into a divergent conformation—one of which ushering in the emergence of the phenylpropanoid pathway.

## Introduction

Plant terrestrialization, the global conquest of land by the ancestors of modern embryophytes, was one of the most consequential evolutionary transitions in Earth’s history [1–3]. This was not a smooth passage. It exposed photosynthetic organisms to a suite of environmental challenges that differed fundamentally from those experienced in an aquatic setting, including fluctuating hydration, temperature extremes, nutrient heterogeneity, mechanical stress, microbial interactions, and elevated irradiance [4, 5]. Among these stresses, ultraviolet-B (UV-B; 280–315 nm) radiation represents a particularly pervasive selective force.

In water, incident UV radiation is attenuated by the water column, dissolved organic matter, and suspended particles [6, 7]; on land, by contrast, photosynthetic tissues may experience rapid and intense changes in UV exposure. UV-B can damage DNA, proteins, membranes, and the photosynthetic apparatus directly, while also promoting the formation of reactive oxygen species (ROS) [8, 9]. The successful establishment of plants on land therefore required not merely tolerance of UV-B damage, but integrated mechanisms for stress perception, signal transduction, cellular protection, repair, and acclimation [9–11]. In land plants, UV-B responses are coordinated across several biological scales. For example, the UV-B photoreceptor UV RESISTANCE LOCUS 8 (UVR8) initiates signaling pathways that alter gene expression and promote acclimation, while other photoreceptors integrate information on light quantity and quality [10, 12–17]. Downstream, a whole suite of responses is mounted. These range from the activation of DNA-repair pathways [18, 19], maintenance and repair of photosynthetic complexes [20], modulation of growth [21], antioxidant responses [22, 23], and the production of UV-absorbing specialized metabolites [24]. Understanding how this regulatory and metabolic toolkit evolved requires experimentally tractable representatives of the algal lineages most closely related to land plants.

Zygnematophyceae are the sister group to embryophytes [3, 25–29]. They are extremely species rich (falling into five orders) and encompass diverse transitions in body plans [30–32]. Owing to their phylogenetic position, zygnematophyte algae provide crucial comparative information that we need to contextualize with data from land plants for reconstructing the molecular physiology of the earliest embryophytes [4, 33–35]. Members of the Zygnematophyceae inhabit diverse freshwater and semi-terrestrial environments. As such, they are often remarkably resilient to high light, desiccation, temperature variation, and UV radiation; their stress tolerance has been associated with cellular restructuring, accumulation of protective metabolites, changes in photosynthetic physiology, and, in several taxa, the formation of extracellular mucilage or pigmented intracellular compartments [4, 36–44]. Further, also directed excretion of pigments into the mucilage has been reported for a peculiar zygnematophyte called *Serritaenia testaceovaginata* [45, 46]. Owing to major progress in the genomic exploration of zygnematophytes [47–51], a couple of species have become accessible to systems biological approaches [33, 35, 52–54].

Zygnematophytes encode homologues of many components involved in land-plant stress signaling. Importantly, these include homologs of the above described hierarchical response system [55], ranging from the photoreceptors, protein kinases, hormone-associated signaling components, regulators of chloroplast homeostasis, and enzymes related to specialized metabolism [4, 27, 33, 34, 47, 50–53, 56–61]. Building on this information, a recent profiling of the UV-B response of the zygnematophyte *Mesotaenium* revealed that phenol-metabolic enzyme homologs, photophysiological regulators and DNA repair factors respond together with upstream kinase, light-quality and homologs of abscisic acid signaling (ABA) components [53]—albeit the ABA cascade we know from land plants operates different and independent of ABA [60, 62, 63]. *Mesotaenium* produces a chemically diverse set of phenolics that accumulate during short term UV response [53]. However, how they integrate UV-response networks for acclimation—and how their outputs compare with those of embryophytes—remains incompletely resolved.

Phenolic compounds, including flavonoids and coumarins, are key for acclimation to UV [9, 64]; they can screen damaging radiation and, in many cases, contribute to antioxidant defense [65]. In land plants, these compounds are sourced from the phenylpropanoid pathway–best known from vascular plants, where it additionally generates lignin, a major structural polymer associated with mechanical support and water transport [66–70]. Comparative genomic analyses have revealed that streptophyte algae possess homologues of several enzyme families associated with phenylpropanoid metabolism—in several cases phylogenetically grouping as co-orthologs to the respective enzymes known from land plant model systems [57]. Yet the occurrence of homologous enzymes does not necessarily imply the presence of the canonical embryophyte phenylpropanoid pathway. Rather, enzyme families may have diversified, been recruited into distinct biochemical contexts, or generated lineage-specific metabolite repertoires before the emergence of land plants [71, 72]. This makes direct metabolite-level evidence essential for determining which chemical strategies were available to the ancestors of embryophytes and which arose later within terrestrial plant lineages.

Here, we build on *M. endlicherianum* as a powerful genome-sequenced system [52, 58, 60, 73] to investigate the response to long-term UV-B exposure using an integrated experimental framework that combines microscopy-based phenotyping and comprehensive transcriptomics and untargeted metabolomics building on large numbers of samples and replicates. We find that long-term UV-B exposure induces the production of several phenolics and the expression of genes involved mainly in DNA repair, cell wall and membrane rearrangement, but also for signature enzymes from the phenylpropanoid pathway. Our data shed light on conserved strings in the web that forms the UV acclimation network.

## Results and Discussion

### Cellular acclimation to repeated daily UV-B exposure in an algal relative of land plants

Phylogenomic efforts have underpinned that Zygnematophyceae are the closest algal relatives of land plants [3, 26, 28, 30, 31, 74] (Figure 1a) and through genome sequencing several Zygnematophyceae have become accessible for systems biological approaches [34, 47, 48, 50, 51]. We built on the single-celled *Mesotaenium endlicherianum* SAG 12.97 as potent system for inferring the molecular chassis shared between zygnematophytes and land plants [35, 52, 60, 75] (Figure 1a). We cultivated *Mesotaenium endlicherianum* at 80 - 90 µmol photons m^−2^ s^−1^ (at a 16 h / 8h day/night cycle) for 7 days and then transferred them to the experimental set-up for an additional one-day-acclimation (Figure 1b). In that setup, they were subjected either to just 50±10 µmol photons m^−2^ s^−1^ growth light (at a 16 h / 8h day/night cycle) or, the UV specimens, to growth light plus a daily 4 hour dose of 0.6 - 1.2 W m^−2^ of ultraviolet (UV) radiation with a peak in the UV-B spectrum (at 311 nm) for 3 consecutive days (Figure 1c). The UV specimens were allowed to recover for 2 days and samples were taken for all downstream analyses at the same time, where all *Mesotaenium* cells had reached a very mature stage of being 13 days old.

**Figure 1.**
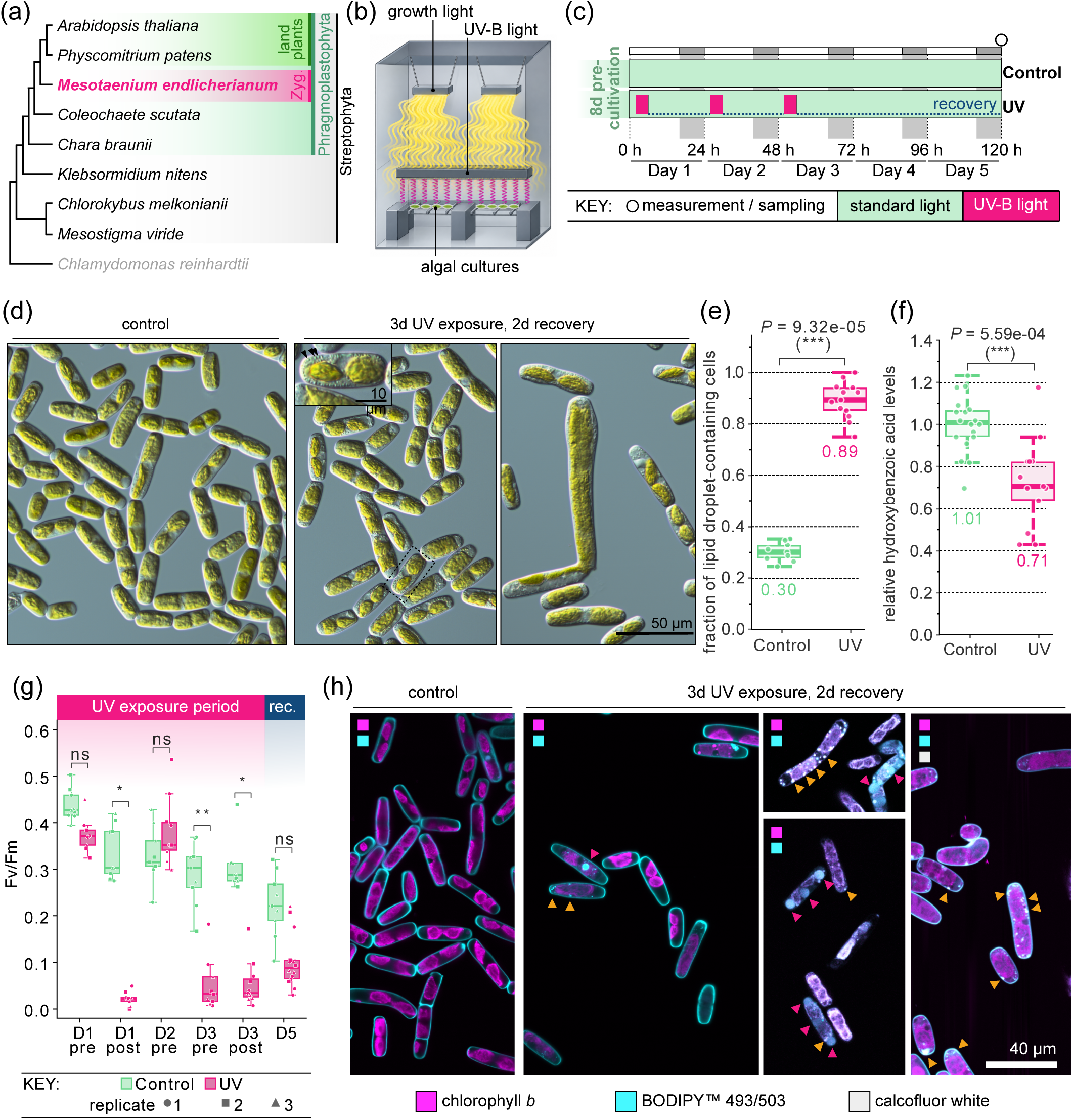
Experimental set-up and physiological response of *Mesotaenium endlicherianum* to repeated UV-B exposure. **(a)** Cladogram showing the phylogenetic position of *M. endlicherianum* (Zygnematophyceae, Zyg.) within the Streptophyta, together with representative streptophyte algae and land plants; the chlorophyte *Chlamydomonas reinhardtii* serves as outgroup. **(b)** Schematic of the cultivation set-up, with algal cultures illuminated by growth light and UV-B light sources; the visual quality of the drawing has bene enhanced using AI. **(c)** Experimental timeline. After 8 d of pre-cultivation under standard light, control cultures remained under standard light, while UV-treated cultures received a UV-B exposure at the beginning of each of days 1–3, followed by a recovery period under standard light until day 5. Grey bars indicate dark periods; the open circle marks measurement/sampling at 120 h. **(d)** Light micrographs of control cells and cells after 3 d of UV exposure and 2 d of recovery. The inset shows a magnified view of the region outlined by the dashed box—note the high number of small inclusions (arrow); the right panel shows elongated/aberrant cells observed after UV treatment. Scale bars: 50 µm (main panels), 10 µm (inset). **(e)** Fraction of lipid droplet-containing cells in control and UV-treated cultures. **(f)** Relative hydroxybenzoic acid levels in control and UV-treated cultures. In (e) and (f), points represent individual measurements, numbers indicate mean values, and *P* values were determined by Mann–Whitney U statistics; \*\*\**P* < 0.001. **(g)** Maximum quantum yield of photosystem II (Fv/Fm) in control and UV-treated cultures, measured before (pre) and after (post) UV exposure on days 1 and 3, before exposure on day 2, and after recovery on day 5. Symbols indicate biological replicates 1–3. Asterisks indicate significant differences between treatments (Welch test; *P < 0.05, **P < 0.01; ns, not significant). (h) Confocal laser scanning micrographs of control cells and cells after 3 d of UV exposure and 2 d of recovery, showing chlorophyll autofluorescence (magenta), neutral lipids stained with BODIPY™ 493/503 (cyan) and cell walls stained with calcofluor white (grey). Coloured squares in each panel indicate the channels shown. Orange arrowheads indicate small peripheral lipid droplets; magenta arrowheads indicate large lipid bodies. Scale bar: 40 µm.

As expected for a 13-day-old *Mesotaenium* culture, all cells appeared sturdy, with chloroplasts streaked with accumulating starch (Figure 1d). Yet, only in the case of UV-exposed *Mesotaenium* cells did we observe a high number of granular inclusions of different sizes (Figure 1d); the fraction of cells showing inclusions increased significantly (*P* = 9.32e-05, Kruskal-Wallis test) in number by almost three-fold, 0.3 compared to 0.89 (Figure 1e). While the stress marker hydroxybenzoic acid was already declining in recovery (Figure 1f), the gross physiological status assessed by maximum quantum yield of photosystem II showed a different trend Fv/Fm (Figure 1g): during the first UV pulse the cells showed a significant drop by almost 0.4, recovering back to the same level as control until the treatment, they exhibited low Fv/Fm values both during the ongoing UV exposure as well as during recovery (Figure 1g).

To scrutinize the identity of the observed inclusions (Figure 1d, e), we used BODIPY™ 493/503 to stain neutral lipids intracellularly (Figure 1h). We found at least three types of lipid inclusions that clearly stained with BODIPY, large drops (Figure 1h, raspberry arrowheads) and smaller droplets (Figure 1h, orange arrowheads). In severely stressed cells with low chlorophyll autofluorescence, these smaller droplets additionally showed strong fluorescence in other channels; whether this reflects for example cellular degradation products remains obscure. Given the presence of clearly identifiable lipid droplets in *Mesotaenium* [58], we interpret at least a subset of these structures as lipid droplets formed during UV acclimation and recovery.

### Chemodiverse responses

Zygnematophyceae produce a range of phenolic compounds under stress—especially upon exposure to strong light or UV. *Zygnema* strains inhabiting polar regions produce UV-absorbing phenolics that are likely stored in vacuoles near the cell surface, and in one strain their amount rose significantly after UV treatment [42]; this was also observed in young vegetative cells of *Zygnema sp.* under UV-A and UV-A+B [42]. In *Zygogonium ericetorum*, filaments from the dry top layer of the mat turn purple and hold much more phenolics and hydrolysable tannins than the green filaments below [76–78]. Upon closer investigation, the pigment responsible for the characteristic purple colour of the vacuoles was resolved to be a highly branched glucose polymer with ester-linked polyphenols such as gallic acid—the purple colour results from complexing/binding of iron [76, 77]. The most striking examples are the glacier ice algae, which construct a brownish vacuolar pigment derived from purpurogallin [79, 80]. In *Mesotaenium berggrenii*, the main compound is purpurogallin carboxylic acid-6-O-β-D-glucopyranoside [79, 81]. *Ancylonema* species also make purpurogallin, partly in conjugated forms [80, 82]. These pigments protect the chloroplasts and nuclei from damaging visible and UV light [79, 81].

We sought to obtain a comprehensive picture of the phenolic bouquet underpinning the long-term acclimation to UV in *Mesotaenium.* For this, we used high resolution mass spectrometry coupled with ultra-high pressure liquid chromatography (UHPLC-QTOF-MS/MS) to generate untargeted metabolomics data from 18 control and 18 samples obtained from long-term UV exposed cells. We performed an unsupervised principal component analysis (PCA) to access global patterns in the metabolomic data (Figure 2a). The metabolomic profiles showed a clear treatment separation along PC1, which explained 31% of the variance in the data, a separation that stayed consistent over the comparisons to the other principal components (Figure 2b). From a total of 2172 features, detected in positive and negative modes (Figure 2c), we focused on features that differed significantly between UV-treated and control samples (Benjamini-Hochberg adjusted *P* ≤ 0.001) and showed a strong change in abundance (log_2_ fold change). Based on this criteria, 74 features were increased and 44 were decreased following UV treatment (Figure 2d; Supplemental Table S1).

**Figure 2.**
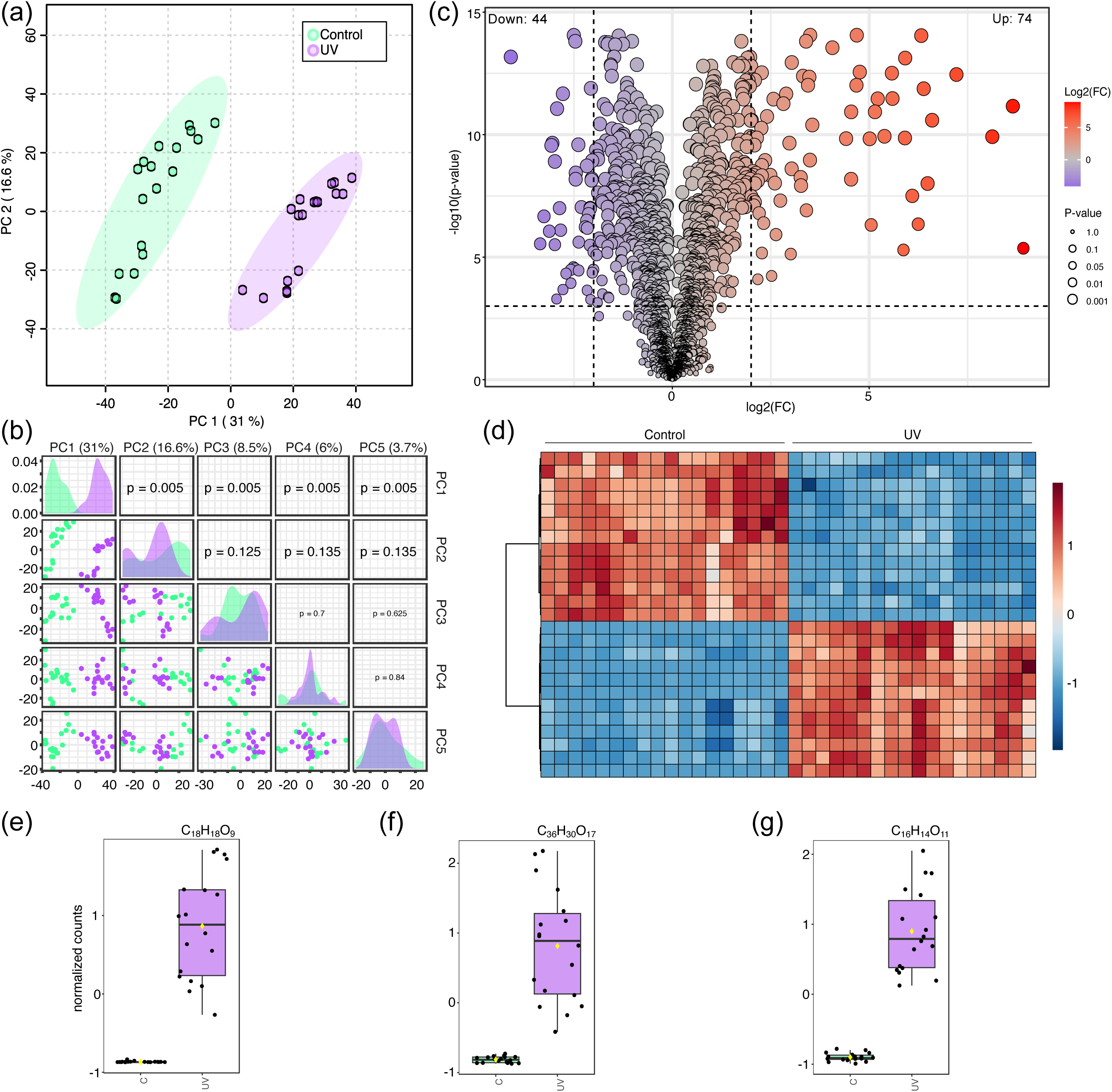
*Mesotaenium endlicherianum* shifts its metabolite fingerprint upon UV acclimation. **(a)** Principal component analysis (PCA) of the untargeted metabolomic profiles of control and UV-treated samples. PC1 and PC2 explain 31.0% and 16.6% of the total variance, respectively. **(b)** Pairwise comparison of the first five principal components (PC1– PC5), showing the distribution of control and UV-treated samples and the corresponding P-values for treatment-associated separation. **(c)** Volcano plot showing differential metabolite feature abundance between UV-treated and control samples. Features with increased and decreased abundance following UV treatment are shown according to log2 fold change and statistical significance (−log10 P-value); 74 features were increased and 44 decreased in their abundance according to the applied significance and fold-change thresholds. **(d)** Heatmap showing the relative abundance patterns of significantly altered metabolite features in control and UV-treated samples. Values are scaled by metabolite feature. **(e,f,g)** Relative abundance of three UV-responsive phenolic metabolites; the molecular formulas calculated based on the exact masses are shown. Points represent individual measurements and boxplots show. Structural drawings represent putative structures/substructures consistent with the MS/MS data.

Our metabolomic analysis revealed several UV-responsive metabolites that accumulated to levels several thousand-fold higher than in the control, highlighting their potential role in UV acclimation (Supplemental tables S1 and S2). Based on the putative annotation by MS/MS, several of these could be phenolic compounds. This could include methoxypsoralen-like hexoside (Figure 2e), putative dimeric hydroxy-methoxypsoralen-like di-hexoside (Figure 2f), and putative polyhydroxycoumarin carboxylic acid O-hexoside (Figure 2g). Methoxyfuranocoumarins such as bergapten (5-Methoxypsoralen, 5-MOP) and xanthotoxin (8-MOP) are linked to UV light in two ways: UV increases their production in plants, and UV activates them photochemically [83–85]. Furanocoumarin production goes up with abiotic triggers such as UV radiation and wounding, and the size of the increase depends on how strong the exposure is. At the gene level, UV raises expression of the prenyltransferase PcPT and increases furanocoumarin output in parsley. In rue (*Ruta graveolens*), UV-B increases expression of CYP98A22, an enzyme that feeds the furanocoumarin pathway [86]. UV can also push these compounds onto the leaf surface, as seen with xanthotoxin in celery, which is thought to protect the plant against UV stress. Overall, methoxyfuranocoumarins seem to act as sunscreens and more as light-activated defences that plants make higher amounts of under UV. Hydroxycoumarin carboxylic acids (a coumarin ring with an OH and a COOH group) are rare as natural products. They occur, for example, in pomegranate [87]. Their several OH groups, especially neighboring pairs, make coumarins good radical scavengers. Synthetic polyhydroxycoumarins confirm this in antioxidant capacity tests like DPPH and ABTS assays and in tests of lipid oxidation. It is conceivable that in *Mesotaenium* under oxidative stress, a hydroxycoumarin carboxylic acid could form from coumarins without any enzyme via hydroxyl radical attack as is the case when generating them synthetically [88, 89]. It could also be a breakdown product rather than something made on purpose—e.g. by breakdown of polyphenolic compounds.

Our data suggest that both short UV treatment [53] and the here presented acclimation to repeated UV induced diverse coumarin-like compounds. The presence of coumarin-like compounds suggests the involvement of the phenylpropanoid pathway. That said, thus far the presence of this biosynthesis pathway in streptophyte algae has remained obscure [53, 90, 91]. And this is despite the fact that homologs likely coding for several key enzymes are present [57], chemodiverse phenolics can be induced under UV in *Mesotaenium* and related species [45, 46, 53, 79], and the induction of derivatives is more pronounced under prolonged exposure in land plants [92] and algae [45]. We thus next turned to understanding the regulation of genes in specialized metabolism.

### Genes in specialized metabolism are among the strongest differentially regulated

To obtain a global view of transcriptional changes associated with acclimation to repeated UV exposure, we performed RNA sequencing in nonaplicates (three biological with three technical replicates each) of RNA from control and UV-treated samples on the illumina platform. We obtained a total of 1,753,186,520 individual reads across 18 samples (237.84 Gbp of data), pseudoaligned the data onto the *Mesotaenium endlicherianum* V2 gene models [58], and calculated global and differential gene expression profiles.

Principal component analysis (PCA) revealed a clear separation between UV-treated and control samples along PC1, which explained 49.4% of the total variance in the data (Figure 3a). The repeated UV exposure induced a pronounced transcriptional response, with a significant upregulation of 451 genes and dowregulation of 444 genes at a (log_2_ fold change of 1 and −1 respectively; BH-corrected *P* ≤ 0.01; Figure 3b; Supplemental Table S3). Our GO term enrichment showed clear signatures of the long-term exposure and repair processes that followed. Down-regulated genes were enriched GO terms speaking of early responses to stimuli, also known from *Mesotaenium* to be fast responding [52, 53, 58, 73]. Up-regulated genes were enriched in repair processes, especially to DNA, and an overall adjustment of the cell physiology, likely facilitated by transporters and cell wall remodelling. Importantly, even from the bird’s eye view, the term “lignin metabolic process” was enriched (Figure 3c)—despite the checkered occurrence of these genes in *Mesotaenium*.

**Figure 3.**
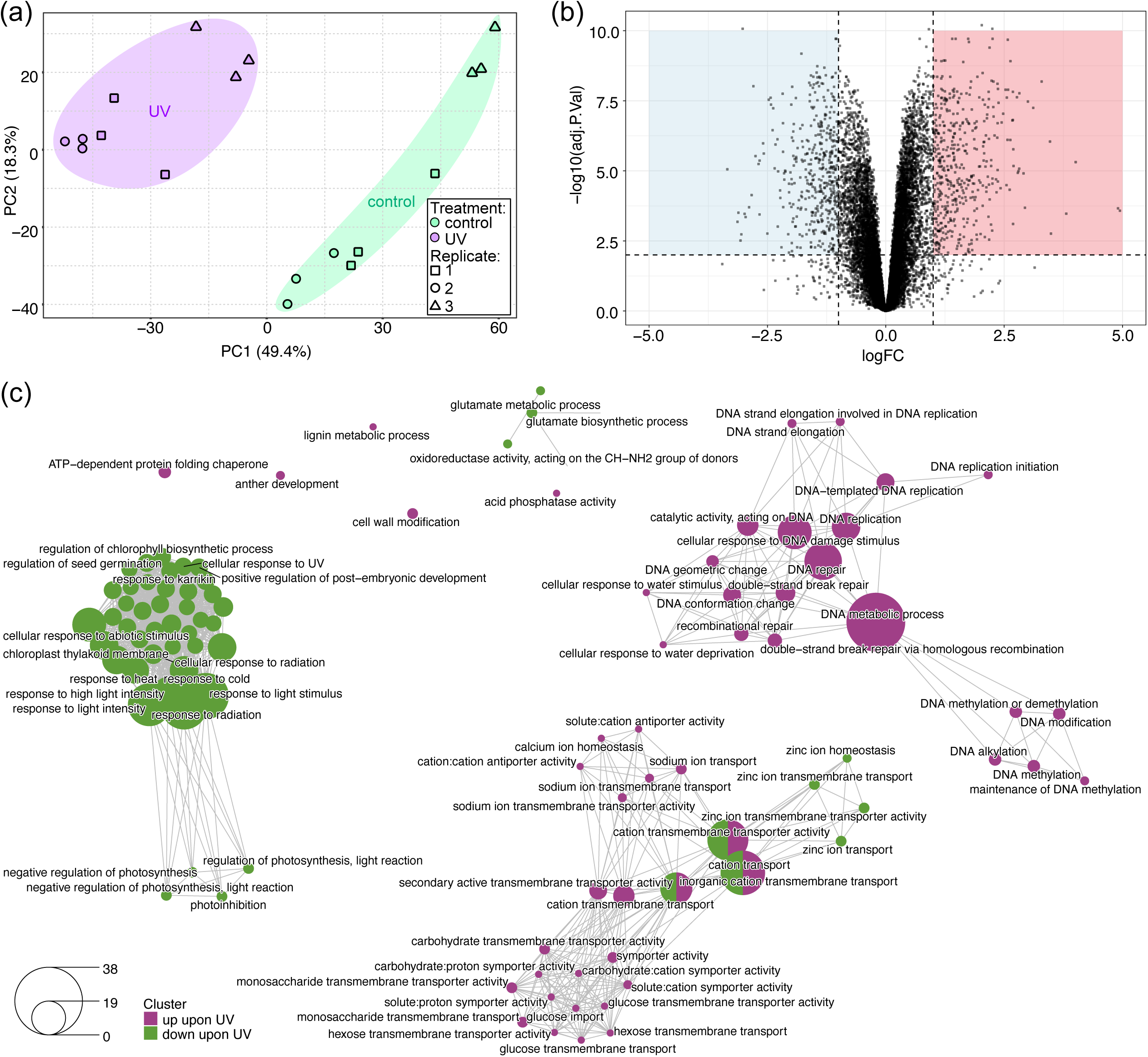
Global gene expression patterns triggered by repeated UV exposure in *Mesotaenium endlicherianum*. **(a)** Principal component analysis (PCA) of the transcriptomic profiles of control and UV-treated samples. PC1 and PC2 explain 49.4% and 18.3% of the total variance, respectively. Colours indicate treatment and symbols indicate biological replicates 1–3. (b) Volcano plot showing differential gene expression between UV-treated and control samples, plotted as log fold change (logFC) against statistical significance (−log10 adjusted P-value). Dashed lines indicate the significance (adjusted P < 0.01) and fold-change (|logFC| > 1) thresholds; shaded areas mark significantly up-regulated (red) and down-regulated (blue) genes. (c) Network of Gene Ontology (GO) terms enriched among differentially expressed genes. Nodes represent enriched GO terms, coloured by whether they are enriched among genes up-regulated (magenta) or down-regulated (green) upon UV exposure; node size corresponds to the number of genes annotated with the term. Edges connect GO terms that share annotated genes, pointing to functional relationships.

Among the top_100_ most differentially up-regulated genes, several salient to UV-induced changes in metabolism and cell biology stand out (Figure 4a and 4b). This includes a glutamine synthetase 1;4 (GS). Cytosolic GS isoforms are often strongly upregulated in response to wounding, heat, and, importantly, UV-B stress — particularly in root tissue [93]. Cytosolic GS1 isoforms (including GLN1;1, the major stress-tolerance contributor) link glutamine biosynthesis to ROS homeostasis in chloroplasts [94]. Among the most upregulated genes (11.2-fold) was a homolog of *ABC TRANSPORTER G FAMILY MEMBER 40* (*ABCG40*), Me1_v2_0234240.1, for which we found no plastid-targeting peptide; data from land plants point to a role of ABCG40 in transporting abscisic acid [95] and potentially other terpenoids [96]—aligning with data from *Serritaenia testaceovaginata* that upregulated a plethora of ABC transporters [46]. Several factors known in cell wall modification were highly up-regulated, including homologs to *CELLULOSE SYNTHASE* 5, *GLYCOSYL HYRDOLASE 9C3*, two pectin lyases, a pectin acetylesterase, glycosyl hydrolases and a *FASCICLIN-LIKE ARABINOGALACTAN 6* (*FLA6*) homolog. FLA proteins likely act in signaling processes in the apoplast [97, 98] and FLA proteins increased in abundance in the proteome of osmotically challenged *Mesotaenium* and *Zygnema* [52].

**Figure 4.**
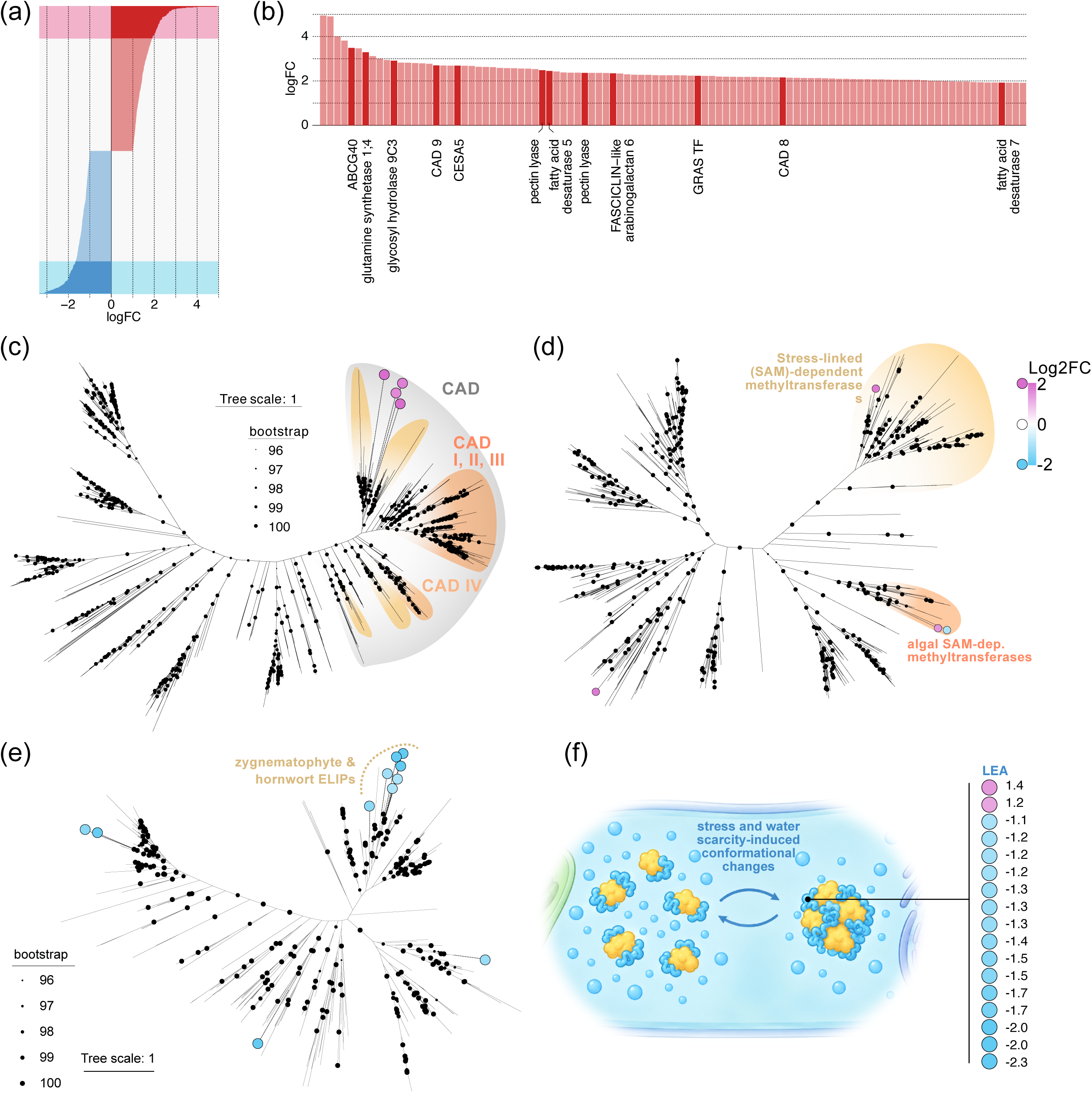
UV-responsive genes and phylogenetic analyses of selected gene families in *Mesotaenium endlicherianum*. (a) Distribution of log fold changes (logFC) of differentially expressed genes upon UV exposure, ranked by logFC. Up-regulated genes are shown in red and down-regulated genes in blue; shaded areas highlight the most strongly up- and down-regulated genes, the up-regulated subset of which is shown in (b). (b) Most strongly up-regulated genes upon UV exposure ranked by logFC; selected genes with annotated functions are highlighted and labelled. (c–e) Maximum likelihood phylogenies of (c) cinnamyl alcohol dehydrogenases (CAD), (d) S-adenosyl methionine (SAM)-dependent methyltransferases and (e) early light-induced proteins (ELIPs). Coloured circles mark differentially expressed *M. endlicherianum* genes, coloured by log2 fold change upon UV exposure (magenta, up-regulated; blue, down-regulated). In (c), the CAD clade (grey) and the CAD I–III and CAD IV subclades are highlighted; in (d), stress-linked SAM-dependent methyltransferases and algal SAM-dependent methyltransferases are highlighted; in (e), the clade of zygnematophyte and hornwort ELIPs is indicated. Black dots on branches indicate UFBOOT support ≥ 96 from 1000 pseudoreplicates, with dot size scaled to support value. Scale bars indicate expected substitutions per site. (f) Schematic illustrating the protective function of late embryogenesis abundant (LEA) proteins, alongside the log2 fold changes of all differentially expressed LEA genes (2 up-regulated, 15 down-regulated); the visual quality of the schematic drawing has bene enhanced using AI.

Our data capture six homologs of cinnamoyl alcohol dehydrogenase (CAD), all of which were upregulated, ranging from 2.1- to 6.5-fold. We mapped them onto the most recent and comprehensive phylogenetic analysis of CAD in zyngematophytes [31] and found that these CADs were all falling into a clade of zygnematophyte CADs branching close to CADI/II/III (Figure 4c). Two additional ones (Me1_v2_0015930 and Me1_v2_0010370) were not captured by the phylogenetic analysis but were nonetheless upregulated. Along the same vein, cinnamoyl CoA reductase 1 (CCR1) was upregulated by 2.5-fold. Two MYB-like genes (homologs of AT2G01060 and RVE1) were upregulated by 2.4- and 2.7-fold. Our data also captured several S-Adenosyl-L-Methionin (SAM)-dependent methyltransferase superfamily proteins; two major groups of SAM-dependent O-methyltransferases act in the phenylpropanoid pathway, Caffeic acid O-methyltransferase (COMT) and Caffeoyl-CoA O-methyltransferase (CCoaOMT). Yet, the differentially regulated (SAM)-dependent methyltransferase showed a different phylogenetic affinity in our maximum-likelihood phylogenetic analysis (Figure 4d): they clustered with putative methyltransferase gene homologs such as AT4G22530 and AT1G55450. The latter has been identified as part of a core stress gene network in Arabidopsis where its activity is regulated under various environmental conditions—including via the ethylene hormonal signaling pathway—as well as during processes involved in the plant immune response (PTI) [99]. Studies on natural variation in Arabidopsis under environmental stress have shown that a loss-of-function mutation in AT4G22530 confers increased resistance to UV radiation in the plants (and also here links to immune response were proposed) [100].

Overall, these upregulations warrant attention. From all data gathered thus far it appears unlikely that the zygnematophytes have a bona fide phenylpropanoid pathway [53, 90, 91, 101–103]. Yet, these data suggest that the homologs that code for enzymes of the phenylpropanoid pathway in land plants have a deeply conserved function in UV responses—irrespective on the compounds on which they act. Studies on other streptophyte algal lineages can illuminate the probability that these enzyme families had UV-responsive roles before the canonical land-plant pathway was assembled.

### Global differential gene expression capture signatures of long-term acclimation instead of rapid light response

Our data capture that cells that have been subjected to UV relatively tune down the fast and early response mechanism to irradiance. This included eleven homologs of EARLY LIGHT-INDUCIBLE PROTEINS (ELIP) that were significantly downregulated from 2.0- to 10.2-fold. EARLY LIGHT-INDUCIBLE PROTEINS (ELIPs) are a family of low-molecular-mass, chlorophyll-binding proteins localized in the thylakoid membranes of plant chloroplasts that protect the photosynthetic apparatus from photooxidative stress caused by oxidative stressors; they achieve this by transiently binding free chlorophyll molecules released during pigment turnover, thereby preventing the formation of harmful singlet oxygen and free radicals [104]. This early transcriptional responses of ELIPs are also a recurrent pattern in zygnematophytes exposed to oxidative stressors [52, 58, 73]. We computed a maximum-likelihood phylogeny of *ELIP* (Figure 4e); while the phylogeny had overall low backbone support for the deep branches, most downregulated *ELIP*s grouped with other zygnematophyte and hornwort sequences.

Along these lines, *GOLDEN2-LIKE 1* (*GLK1*), a transcription factor that acts as master regulator of photomorphogenesis and photosynthesis-associated gene expression [105–107], was also downregulated. This is noteworthy because GLK1, and its interconnectivity in gene expression data, is a synapomorphy of zygnematophytes and land plants [58]. Similarly, *CAROTENOID CLEAVAGE DIOXYGENASE 1* (*CCD1*) was downregulated. Stress-responsive regulation of *CCD1*-like genes has previously been observed in zygnematophytes, where carotenoid turnover and apocarotenoid formation form part of the oxidative-stress response [73, 108]. Its downregulation here may therefore reflect the transition from an acute carotenoid-derived stress response toward a longer-term acclimated state. Indeed, carbon fixation even could be ramping up, since a homolog of the α-carbonic anhydrase CAH3, Me1_v2_0123140, was upregulated 2.2-fold (P = 1.4e-3).

Following suit, other stress marker genes such as *LATE EMBRYOGENESIS ABUNDANT* and *EARLY RESPONE TO DROUGHT* were downregulated. For example, 15 homologs of *LEA* were downregulated 2.1 to 5.0-fold while only two were upregulated (2.7- and 2.3-fold; Figure 4f). Because transcripts for these genes are depleted in comparison to the control it suggests a global re-routing of the cell acclimation towards a different mode of action: cells enter a new specialized metabolic-based homeostasis. It is further possible that this is part of a feedback loop that—after strong increase in transcripts for these genes then undergoes transcriptional attenuation upon cessation of the stressor, as expression levels fall below the pre-stress baseline, indicative of a state of adaptive quiescence distinct from the unstressed condition.

### Are plant phenolics born out of UV response?

In vascular plants, phenylpropanoid metabolism can represent a major sink for fixed carbon, particularly through the production of lignin—one of the most abundant biopolymers on earth [109]. Photosynthetic organisms constantly fix CO₂ and must channel it somewhere, and carbon-concentrating mechanisms across the green lineage have probably made carbon relatively overabundant. This low cost might have given streptophytes room to experiment with carbon-rich specialized metabolites, tolerate some metabolic “noise”, and keep new compounds that were neutral or only slightly beneficial [72]. Our data suggest that the enzyme toolkit of the phenylpropanoid pathway has a conserved role in UV protection. This raises the exciting question of whether the pathway’s deep evolutionary origin is tied to UV response.

Phenylpropanoid-derived compounds such as flavonoids are induced by UV in flowering plants, in the moss *Physcomitrium patens* and in the liverwort *Marchantia polymorpha* [64, 92, 110, 111]. We here found a set of compounds in *Mesotaenium* that one would tie to necessitating an active phenylpropanoid biosynthetic pathway, aligning with: (a) flavonoids that were detected in the zygnematophyte alga Penium margaritaceum [48]; (b) zygnematophyceae that produce further phenolic UV screens such as purpurogallins [79, 80]. Overall, this suggests that shikimate- and phenylpropanoid-derived phenolics have deep roots. Yet genes coding for canonical enzymes, such as PHENYLALANINE AMMONIA LYASE (PAL) and CINNAMATE 4-HYDROXYLASE (C4H) candidates are largely missing from streptophyte algal genomes [57, 90, 112], even though *p*-coumaric acid occurs in diverse green algae [113], a *PAL* homolog occurs in Klebsormidiales [57, 112, 114], and a divergent PAL acts in *Chara* [115]. This implies non-canonical routes to phenylpropanoid building blocks. Indeed, the evolution of the phenylpropanoid pathway in streptophytes is not only rich in expansions [57] but also loss and contraction [116], showcased by the reduced set of phenylpropanoid biosynthesis-related genes in the purpurogallin-producing ice alga *Ancylonema nordenskiöldii* [91, 117]. It further implies that the non-canonical routes were assembled from deep homologs of phenylpropanoid enzymes and promiscuous enzyme families such as cytochrome P450s.

Regulation may be even more conserved than biosynthesis. A multilayered molecular network operates in fine-tuning the response of embryophytes to environmental challenge that ranges from perception of a cue, to signal transduction, to regulation of transcription, and metabolic outputs [55]. At each layer, multiple regulatory players operate with both overlapping and specific functions. This means that various routes can lead to similar outcomes but also several branching points along the hierarchy can occur. MYB transcription factors are well known for their role in regulating the accumulation of phenolics upon UV stress, whose action ultimately results in the elevated production of flavonoids and anthocyanins [118–120]. In *Marchantia*, the UV-B-responsive MYB14 module controls auronidins, which are anthocyanin-like pigments made from aurones [121] that however only occur in liverworts. This suggests that the regulatory logic of UV protection was conserved while the protective metabolites themselves varied between lineages [72]. Similarly, the NAC–MYB network that directs lignification in vascular plants also operates in non-lignified moss hydroids and includes MYBs related to UV-B-induced moss MYBs [122]. The network is probably at least 500 million years old and could have acted on phenylpropanoid metabolism before lignin even existed.

Together, these observations suggest a stepwise scenario. First, UV pressure selected for the formation and hyperaccumulation of phenolic compounds, many of which were deposited in cell walls and surface structures. The cuticle of *P. patens* is enriched in UV-shielding phenylpropanoid-derived phenolics [123]. The UV-tolerant Antarctic moss *Ceratodon purpureus* holds about nine times more UV-shielding phenolics in its cell wall than in its soluble fraction [124]. Coumarates and flavonoids are also built into sporopollenin, a UV shield around spores and pollen. Aromatic, sporopollenin-like polymers (algaenan) around the spores of filamentous zygnematophytes like *Mougeotia*, *Zygnema*, and *Spirogyra* hint at a similar role in algae [44, 125–130]. Once these phenolics were accumulating in large amounts and being deposited into walls, the step to polymerisation was short. The resulting lignin and lignin-like substances brought new structural and water-conducting advantages on land. Under this rationale, lignin is an exaptation: an expanded, UV-driven phenolic metabolism that was later co-opted as the main carbon sink of vascular plants[72].

## Conclusion

UV is a prime terrestrial stressor. We here used a setup where *Mesotaenium* had to meet this challenge with a stable, metabolism-based acclimation rather than a transient emergency response (Figure 5). This suggests the capacity for lasting UV acclimation was already present before plants colonized land. A response that builds on phenolics. The consistent recruitment of phenylpropanoid enzyme homologs, even without a canonical pathway, suggests that phenylpropanoid metabolism did not arise as a single complete pathway. Instead, it was probably assembled step by step from an older, UV-responsive enzyme toolkit whose substrates and products changed over evolutionary time (Figure 5). If so, the evolution of lignin and other wall-bound phenolics looks less like a sudden innovation of vascular plants. It looks more like the co-option of an ancient protective chemistry that let embryophytes turn surplus carbon into sunscreens and, eventually, into structural polymers. While the molecular links remain obscure, both phenolics and enzyme-coding homologs that act in embryophytic phenylpropanoid biosynthesis constitute a shared set of the UV response toolkit of the last common ancestor of land plants and algae.

**Figure 5.**
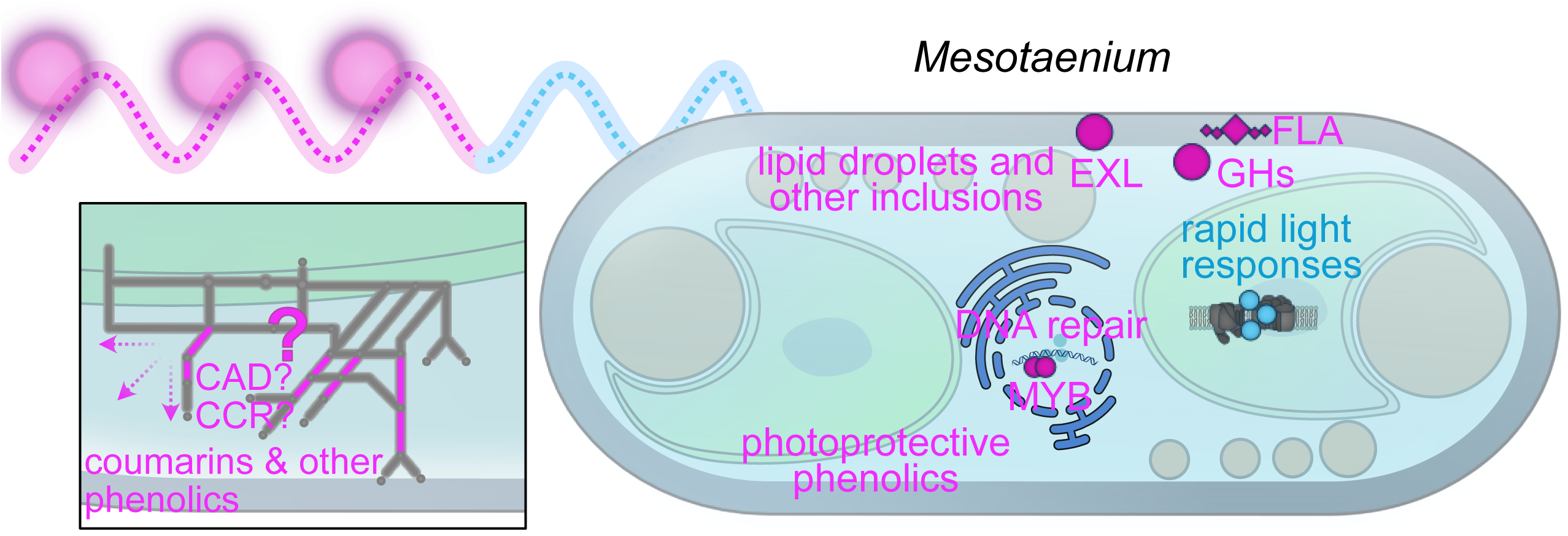
Summary model of the response of *Mesotaenium endlicherianum* to repeated UV-B exposure. Schematic drawing of a *M. endlicherianum* cell summarising the processes affected by UV-B (magenta wave) followed by ongoing during recovery (blue wave). UV-B exposure is associated with accumulation of lipid droplets, induction of DNA repair, changes in MYB transcription factor and gene expression of cell wall modifiers, and the production of photoprotective phenolics; rapid light responses such as *EARLY LIGHT-INDUCED PROTEINs* are downregulated. The inset illustrates the the land-plant phenylpropanoid network as reference, with steps for which UV-responsive Mesotaenium homologs exist highlighted in magenta highlighting the patchiness; the question mark indicates the overall unresolved biosynthetic routes.

## Acknowledgement

We thank Prof. T. Friedl from the Department of Experimental Phycology and Culture Collection of Algae (SAG), Prof. S. Neugart and René Heise at the University of Göttingen for their support. J.d.V. is grateful for funding by the German Research Foundation (DFG) grants 509535047 (VR 132/10-1) and 569599232 (VR 132/18-1) and the grants 440231723 (VR 132/4-1), 528076711 (VR 132/13-1) within the framework of the Priority Programme “MAdLand – Molecular Adaptation to Land: Plant Evolution to Change” (SPP 2237). J.d.V. further thanks the European Research Council for funding under the European Union’s Horizon 2020 research and innovation programme (Grant Agreement No. 852725; ERC-StG “TerreStriAL”) and the Horizon Europe programme (Grant Agreement No. 101230161; ERC-CoG “StreptoProgram”). I.F. acknowledges funding by the DFG grants 495720893 (INST 186/1434-1) and 440232164 (FE 446/14-1) within the framework of the Priority Programme “MAdLand – Molecular Adaptation to Land: Plant Evolution to Change” (SPP 2237). C.F.K is grateful for being supported through the International Max Planck Research School (IMPRS) for Genome Science within the framework of the ‘Göttingen Graduate Center for Neurosciences, Biophysics, and Molecular Biosciences’ (GGNB) at the University of Goettingen. C.F.K. thanks the IMPRS Genome Science and the Rahn-Quade Foundation (Rahn-Quade-Stiftung Förderung zusätzliche Sachmittel T0381/49228/26).

## Methods

### Data availability

All RNAseq data have been deposited on NCBI under Bioproject PRJNA1504534

### Algal strains

*Mesotaenium endlicherianum* SAG 12.97 was obtained from the Department Experimental Phycology and Culture Collection of Algae at Göttingen University (EPSAG) [131].

### Cultivation of Algae on Solid Medium

For stock culturing, *M. endlicherianum* was stored on solid Bold’s Basal Medium (BBM) with vitamins and triple nitrate (3NBBM+ V) according to the recipe of Schlösser et al. [132] (see also Bischoff and Bold [133] and Starr and Zeikus [134]) at 15 °C under full-spectrum fluorescent lamps (5-10 μmol photons m^−2^ s^−1^; constant light). Stock cultures between 21 and 28 days were used for the inoculation of experimental plates.

For easier sampling and reduction of agar and media contamination in metabolite samples, the agar was covered with cellophane (folia®, Max Bringmann KG). The cellophane was cut to petri dish size, autoclaved and applied using a heated Drigalsky spatula and five drops of sterile ddH_2_O. The biomass of one stock plate was diluted in 10 mL liquid medium at room temperature and 300 μL of the solution was spread on each plate using a sterile cooled Drigalsky spatula. Plating occurred under the sterile bench with sterile technique. The plates were closed using 3M Medical Micropore Tape, as it was shown to ensure better CO_2_ exchange than parafilm[135].

For the UV-B exposure experiments, the plates were grown at 80 − 90 μmol photons m^−2^ s^−1^ (Niello® LED 300W, 380–740 nm spectrum) for seven days with a 16/8 h light/dark cycle, before being transferred to the UV setup location for 24 h acclimation before the experiments.

### Experimental Set-Up

The UV setup used the same grow lights as previously described in addition to two UV-B lamps used in conjunction, the UV-B Broadband TL fluorescent tube lamp with a UV-B spectrum of 290 to 315 nm, and the UV-B Narrowband TL fluorescent tube lamp with a UV-B spectrum between 305 and 315 nm, which peaks at 311 nm (lamp wattage: 40 W and 20 W; length: 1213.6 mm and 604 mm respectively, Philips, the Netherlands; spectrum Supplementary Fig. S1A). They produced a UV-B intensity ranging from 0.6 – 1.2 W m^−2^, depending on location of the individual petri dish. The photosynthetically active radiation (PAR) values were 40 − 50 μmol photons m^−2^ s^−1^and 50 − 60 μmol photons m^−2^ s^−1^ for UV-B and control, respectively. While PAR was applied with the previous 16/8 h light/dark cycle, for UV treated samples UVR was applied for 4 h 3 d in a row (Fig. 1b, 1c) To reduce condensation on the lids of the petri dishes, fans were installed to create air circulation. Temperature and humidity were monitored using a Tempo Disc™ (BlueMaestro). The temperature and humidity ranged from 19.7 °C to 24.3 °C with an average of 22.3 °C and 41.1 % to 68.6 % with an average of 51.0 % for control and 19.6 °C to 26.5 °C with an average of 23.1 °C and 42.7 % to 73.2 % with an average of 54.3 % for UV, respectively.

For the multiple day experiment, the petri dish plastic lids were left on, which resulted in a reduced UV-B intensity of 0.6 – 1.2 W m^−2^. The measurements of photosystem II (PSII) operating efficiency (Fv/Fm) were taken from three places on three independent plates using a MINI-PAM II (Walz, Germany). For statistics, technical replicates were averaged and UV versus control were compared with a Welch’s t-test [136] and *P* values were adjusted using Holm’s method [137].

Morphology assessment through microscopy was performed at 120 h using the Olympus BX-60 microscope (Olympus, Japan) with DIC equipped with JENOPTIK GRYPHAX® PROKYON camera and JENOPTIK GRYPHAX® microscope camera software (version V2.2.0) (JENOPTIK AG, Jena, Germany). The imaging occurred immediately after measurement. Additional confocal images were taken at the same sampling timepoint. The cells were stained using 1% Calcofluor White Stain (Sigma-Aldrich, Germany), Invitrogen™ pHrodo™ Green AM Intracellular pH Indicator Dye and Invitrogen™ BODIPY™ 493/503. The images were taken using the Zeiss LSM 980 Confocal Laser Scanning Microscope equipped with diode lasers 405nm, 445, 488, 514nm; diode-pumped solid state lasers 543, 561, 594, 639nm all with the objectives 40x/1,4Plan-Apochromat and an Axioxam 512 color camera using Zeiss Zen blue software V3.2 (Zeiss, Jena, Germany). The fraction of cells containing lipid droplets was quantified per field of view for control (C, nine field of views, 664 cells) and UV-treated (UV, thirteen field of views, 584 cells) conditions. As the data were not assumed to follow a normal distribution, differences between groups were assessed using the Kruskal-Wallis H test [138], a non-parametric alternative to one-way ANOVA. Statistical analyses were performed in Python (v3.12.7) using the scipy.stats module (v1.13.1). A significance threshold of α = 0.05 was applied; p-values are reported alongside the H statistic. Data are presented as boxplots showing the median, interquartile range, and individual data points.

Sampling was performed as follows: six plates each at the time point of the experiment were taken out of both settings, *Mesotaenium* was removed from the cellophane and 1/4 of two plates pooled (depending on setting) for transcriptomics and 3/4 of each plate for metabolite analysis. Pooling was done to balance the varying irradiation intensity. Samples were frozen immediately using liquid N_2_ and stored at −70 °C, to stop any (non-) enzymatic reaction which would distort the results. The experiments were repeated three times to obtain biological triplicates, and each biological replicate had three and six technical replicates for transcriptomics and metabolomics, respectively.

### RNA isolation and sequencing

The Sigma Aldrich Spectrum™ Plant Total RNA-Kit was used for RNA extraction, following the “Spectrum Plant total RNA Isolation protocol A” protocol with the following modifications. 500 µL of the prepared lysis solution/2-mercaptoethanol mixture was added to the sample along with 1/3 beads from MPI Lysing Matrix E. This was immediately vortexed and the biomass homogenized using a Biospec Mini-BeadBeater for 30 s at 5000 rpm. Afterwards, 200 µL lysis solution/2-mercaptoethanol mixture was added, samples placed in an ultrasonic bath (DK Sonic DK-300H, 40 KHz, 120 W) for 2 min with constant ultrasonication. After incubation at 56 °C for 5 min, the solution was transferred to a new Eppendorf tube, avoiding the BeadBeater beads and centrifuged for 1 min at 16 000 x g to remove cell debris and remaining beads. The supernatant was loaded onto the filtration columns. To bind RNA, protocol A with increased amount of Binding Solution of 750 µL was used and subsequently, the manufacturers protocol was followed. RNA concentration and quality was assessed using a NanoDrop Spectrophotometer ND-1000 and 1% agarose gel. 1 µL 6 x loading dye was added to 4 µL sample and incubated at 60 °C and on ice for 5 min each before loading onto the gel. The remaining sample was treated with DNAse I according to manufacturer’s protocol (Thermo Fisher). The total RNA samples used for transcriptome analyses were again quantified, this time using a Qubit RNA HS Assay (Thermo Fisher Scientific, MA, USA). Quality was assessed by capillary electrophoresis using the Fragment Analyzer and the ‘Total RNA Standard Sensitivity Assay’ (Agilent Technologies, Inc. Santa Clara, CA, USA). Library preparation was performed using the ‘Illumina stranded mRNA Prep, Ligation’ kit following the manufacturer’s instructions (Illumina Inc. San Diego, CA, USA).

Briefly, to enrich the mRNAs from total RNA,a polyA-capture was used, followed by fragmentation, the reverse synthesis of cDNA, adapter ligation, and library amplification. Bead purified libraries were normalized and finally sequenced on the NovaSeqX Plus platform (Illumina Inc. San Diego, CA, USA) set to 2×151 bp. The bclconvert tool (version 4.3.6) was used to convert the bcl files to fastq files as well for adapter trimming (Read 1 and Read 2 adapter trimming: ACTGTCTCTTATACACATCT) and demultiplexing. We sequenced nonaplicates (three biological with three technical replicates each) of RNA from control and UV-treated samples on the illumina platform. We obtained a total of 1,753,186,520 individual reads across the 18 samples (237.84 Gbp of data)

### Transcriptome analysis

The quality of the raw and processed reads was checked using FastQC (version 0.12.1) [139] and MultiQC (version 1.16) [140]. Trimmomatic (version 0.39) [141] was used to process raw reads with the setting “ILLUMINACLIP:adapters/adapter.fasta:2:30:10:2:True LEADING:26 TRAILING:26 SLIDINGWINDOW:4:20 MINLEN:36”. The processed reads were pseudoaligned to cDNA of *M. endlicherianum* SAG 12.97 gene model V2 [58] with Kallisto (version 0.48.0) [142] with 100 bootstraps. Using kallisto [142], we pseudoaligned the reads onto the V2 gene models[58] of the *Mesotaenium* genome [47]. The samples were sequenced with the rf-stranded technique. Subsequent analysis was performed using R version 4.6.1. The transcript-level estimates were imported via the TxImport package (v. 1.40.0) [143] and estimated counts calculated based on abundance with “lengthScaledTPM”. edgeR (v. 4.10.5) [144] was used to filter (omitted genes with less than 1 count per million in at least 3 samples) and normalise (upperquartile method [145]) the counts. The data was subsequently transformed to log2-count per million via the voom function in the limma package (v3.68.5) [146, 147]. The principal component analysis was performed with stats (v. 4.6.1) and visualised with ggplot2 (v. 4.0.3) [148]. Functional annotation of genes via eggnog-mapper retrieved from Dadras et al. 2023 a. Gene ontology (GO) over-representation analysis (ORA) was performed using the clusterProfiler enricher function (v. 4.20.0) [149] using a q-value and adjusted p-value cutoff of 0.05 each. The p-values were calculated using the Student test (Student [150]) and adjusted with the Benjamini and Hochberg (BH; Benjamini & Hochberg [151]) method.

### Sample Preparation and Metabolite Extraction

Cultures were transferred into Eppendorf tubes and lyophilized overnight. Two tungsten beads were added to each tube, and the material was pulverized using a Mixer Ball Mill MM400 (Retsch). Approximately 5 mg of dried and homogenized algal material was extracted with 500 μL of 70% (v/v) methanol, following the protocol described by Kelly et al.[152]. After centrifugation, 200 μL of the supernatant was transferred to an LC vial and dried using a SpeedVac. Prior to analysis, samples were reconstituted in 20 μL methanol and subsequently diluted with 60 μL Milli-Q H₂O.

### Untargeted Metabolomics Analysis of Mesotaenium Cells

Samples were analyzed using an ultra-high-performance liquid chromatography (UHPLC) system (1290 Infinity, Agilent Technologies, Santa Clara, CA, USA) coupled to a quadrupole time-of-flight mass spectrometer (QTOF-MS; 6546 UHD Accurate-Mass QTOF, Agilent Technologies). Data were acquired in positive and negative electrospray ionization modes using chromatographic and mass spectrometric conditions as described in Kunz et al. (2026). Raw MS data were processed using MassHunter Profinder B.08.00 (Agilent Technologies) employing recursive feature extraction, as described by [153]. Feature intensities were normalized to sample dry weight, and statistical analyses were performed using MetaboAnalyst 6.0 [154].

Metabolite annotation was performed according to the workflow established in Kunz et al. [53], using accurate mass, retention time, and MS/MS fragmentation information and comparison against the in-house and *Mesotaenium*-specific metabolite databases described therein.

### Phylogenetic analyses

To understand the evolutionary history of FAD and ELIPs, we used manually selected homologs as queries in a BLASTp search against a protein database inferred from the genomes of: *Arabidopsis thaliana* (Lamesch et al. [155]), *Arabidopsis lyrata* (V2.1; [156, 157]), *Arachis hypogaea* [158], *Brassica oleracea* [159], *Brassica rapa* [160] (FPsc v1.3;), *Capsella grandiflora* [161], *Triticum aestivum* [162], *Carica papaya* [163], *Oryza sativa* [164] (v7), *Brachipodium distachyon* [165] (v3.2), *Amborella trichopoda* [166], *Gossypium hirsutum* [167], *Theobroma cacao* [168], *Solanum lycopersicum* [169], *Nicotiana tabacum* [170]*, Zea mays* [171] (V4), *Gnetum montanum* [172], *Glycine max* [173], *Picea abies* [174], *Azolla filiculoides* [175], *Isoetes taiwaniensis* [176], *Selaginella moellendorffii* [177], *Physcomitrium patens* [178], *Sphagnum fallax* [179], *Marchantia polymorphya* [180], *Anthoceros punctatus* [181], *Anthoceros agrestis* [181], leveraging the diverse streptophyte algal genomes[34] including, *Mesotaenium endlicherianum* [47] (building on the v2 gene models [58]), *Penium margaritaceum* [48], *Spirogloea muscicola* [47], *Chara braunii* [182], *Klebsormidium nitens* [183] (v1.1), *Chlorokybus melkonianii* [184] — for naming, see Irisarri et al. [185], *Mesostigma viride* [184], *Zygnema circumcarinatum* SAG 698-1b, UTEX 1559 and UTEX 1560 as well as *Zygnema cylindricum* SAG 698-1a genome [50], *Spirogyra pratensis* [51] (strain MZCH10213), *Micromonas pusilla* [186] (v3.0), *Ostreococcus lucimarinus* [187] (v2.0;), *Ulva mutabilis* [188], *Chlamydomonas reinhardtii* [189] (v5.6), and *Chlorella variabilis* (Blanc et al. [190]). All significant hits (*e* value cutoff of 10^−5^) were aligned with MAFFT [191] (v7.490; L-INS-I). We computed maximum likelihood trees using IQ-TREE multicore version 2.0.7 for Linux 64-bit [192] with 1000 ultrafast bootstrap pseudo-replicates [193]. The best models for protein evolution were determined by ModelFinder [194] and the best model according to Bayesian Information Criterion was used.

